# HOT or not: examining the basis of high-occupancy target regions

**DOI:** 10.1101/107680

**Authors:** Katarzyna Wreczycka, Vedran Franke, Bora Uyar, Ricardo Wurmus, Altuna Akalin

## Abstract

High-occupancy target (HOT) regions are the segments of the genome with unusually high number of transcription factor binding sites. These regions are observed in multiple species and thought to have biological importance due to high transcription factor occupancy. Furthermore, they coincide with house-keeping gene promoters and the associated genes are stably expressed across multiple cell types. Despite these features, HOT regions are solemnly defined using ChIP-seq experiments and shown to lack canonical motifs for transcription factors that are thought to be bound there. Although, ChIP-seq experiments are the golden standard for finding genome-wide binding sites of a protein, they are not noise free. Here, we show that HOT regions are likely to be ChIP-seq artifacts and they are similar to previously proposed “hyper-ChIPable” regions. Using ChIP-seq data sets for knocked-out transcription factors, we demonstrate presence of false positive signals on HOT regions. We observe sequence characteristics and genomic features that are discriminatory of HOT regions, such as GC/CpG-rich k-mers and enrichment of RNA-DNA hybrids (R-loops) and DNA tertiary structures (G-quadruplex DNA). The artificial ChIP-seq enrichment on HOT regions could be associated to these discriminatory features. Furthermore, we propose strategies to deal with such artifacts for the future ChIP-seq studies.

## Introduction

Chromatin immunoprecipitation followed by sequencing (ChIP-seq) is now a standard method to quantitatively assay the binding sites of a DNA binding protein in the genome. Large scale projects such as ENCODE [1] and modENCODE [2] used this technology to find the binding sites of hundreds of proteins in multiple species. With more binding site data available, it has become apparent that certain parts of the genome harbour high frequency of protein-DNA binding events. These regions are called high-occupancy target (HOT) regions and they are observed in multiple species [3,4]. HOT regions are associated with housekeeping genes and are enriched in binding events without canonical motifs [5]. HOT regions are thought to have biological importance due to high number of binding sites observed, but previous reports failed to assign a clearly distinctive function that would explain the requirement for the exuberant number of bound transcription factors.

In this study, we aim to gain a deeper understanding of the nature of HOT regions and the genomic features associated to them. First, we wanted to investigate the features that are common to HOT regions across species. To date, there has been no cross-species comparison of HOT regions in terms of sequence features. The sequence features that are shared across species can lead to a better understanding and prediction of HOT regions in other species as well. With the sequence analysis and subsequent integrative analysis, we primarily aim to uncover the rationale behind the propensity of HOT regions to have unusual number of binding events, many of which are motifless binding events (transcription factors binding to a region without the known motif) [5]. For us, the plausible explanations for motifless binding are a combination of 1) interaction of transcriptions factors (TFs) where only a handful of them are actually binding to DNA [3], 2) existence of weak binding sites where TFs bind to non-canonical motifs in a weak manner [6], 3) regions with high-affinity for chromatin immunoprecipitation called “hyper-ChIPable” regions [7]. Many of the HOT regions are shown to bind hundreds of proteins based on ChIP-seq experiments [4]. However, the stable interaction of a protein complex that consists of hundreds of TFs or side-by-side binding of TFs, many to weak motifs, seems implausible. Detection of hundreds of proteins occupying an individual hot region could be explained by extensive protein interaction networks between transcription factors and cofactors, where only a few factors directly bind to DNA. However, only a handful of such interactions were experimentally validated [3]. Side-by-side binding of hundreds of proteins is again not possible due to space limitations, because the average size of a transcription factor is 10 nm [8] and if a hundred of them bound next to each other as packed as possible, it would require 1000 nm in length (or about ~2900 bp assuming 1bp is 0.34nm [9]), which is beyond the size of average HOT regions. Therefore, we seek additional explanations for existence of HOT regions in the genome and their association with motifless binding.

For a better understanding of what creates the HOT regions, firstly, we investigated nucleotide sequence features (motifs, k-mer content, etc.) of HOT regions across the species. We built species-specific machine-learning models to learn discriminative sequence features for HOT regions. We showed that HOT regions are associated with certain sequence features that are shared across species. Secondly, in order to investigate the potential technical biases causing the occurrence of HOT regions, we analyzed ChIP-seq experiments for knocked-out transcription factors in mice. Previously, [7] showed that highly expressed loci in yeast give rise to false-positive peaks when they did ChIP-seq experiments for proteins that did not have the corresponding gene in the genome. We set out to examine if such a technical bias could be the driving force behind HOT regions given that motifless binding is prominent on those regions. As a result of this analysis, we observed false positive signals on HOT regions. Finally, we investigated the association of HOT regions with RNA:DNA hybrids called R-loops. The two classes of regions share similarity in sequence features and genes associated. We demonstrated association of HOT regions with R-loops. This paper presents a new rationale that explains the apparent high-occupancy on at least some of the HOT regions. With a better understanding of HOT regions provided here and other potential sources of bias associated with false-positive ChIP-seq peaks, now researchers can avoid these pitfalls and obtain less noisy data by additional computational analysis.

## Results

### HOT regions exist in multiple species and cover TSSes of stably expressed genes across cell types

HOT regions are observed in multiple species - human [3,10], *D. melanogaster* [11], yeast [12] and *C. elegans* [10,13]. Based on density of ChIP-seq peaks used as a measure of TF occupancy, we defined HOT regions in human, mouse, worm (*C. elegans)* and fly (*D. melanogaster*) (See Figure 1A and Methods). Our method detected 4324 HOT regions in human, 2638 in mouse, 422 in *C.elegans* and 408 in *D. melanogaster*, out of 428,498 regions with at least one peak in human, 245,250 in mouse, 40,921 in *C. elegans*, and 37,853 in *D. melanogaster*. These examined regions along with their TF occupancy percentiles (percentiles from ChIP-seq peak count distribution) as well as HOT regions are accessible via UCSC track hub (https://bimsbstatic.mdc-berlin.de/hubs/akalin/HOTRegions/hub.txt).

**Figure 1.**
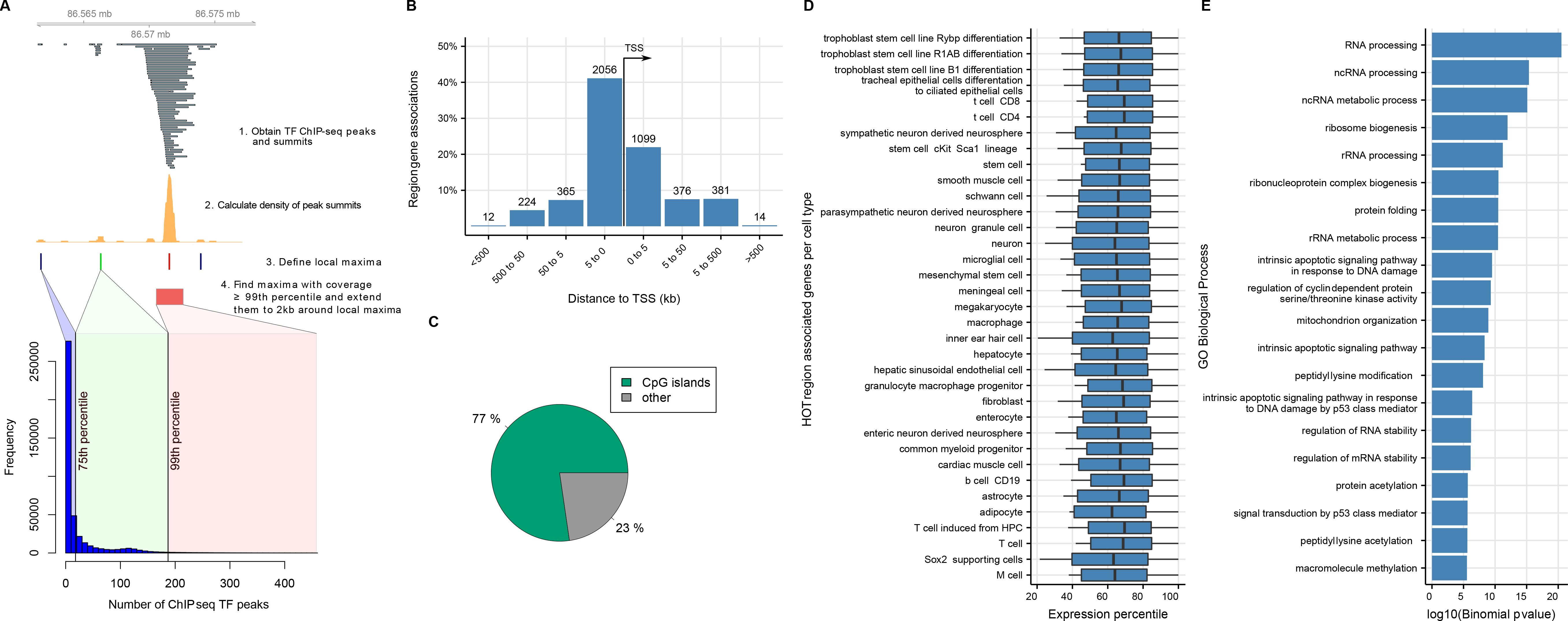
Features of HOT regions. A) Schematic workflow of HOT regions definition. The barplot indicates number of ChIP-seq peaks in HOT (red), MILD (green) and COLD (blue) regions. B) HOT regions are located mostly close to transcription start sites and are promoter associated. The figure shows binned orientation and distance between HOT regions and the nearest genes. Associations precisely at 0 refers to the transcription start site of the nearest gene. C) Most HOT regions overlap with CpG islands. D) Expression profiles of genes associated with HOT regions across cell types in mouse (Expression Atlas EBI databases using fantom5 CAGE expression). The genes are stably expressed in all 35 cell types between 40th and 80th percentile. E) Functional enrichment analysis with Gene Ontology and and KEGG pathway on genes associated with murine HOT regions. HOT regions are significantly enriched for terms that relate to housekeeping functions.

HOT regions are typically located at promoters. The majority of HOT regions (80%) are in close proximity to TSS (within 5 kb) of genes (Figure 1B). In human and mouse, they are mostly associated with CpG islands (Figure 1C). The analysis of properties of genes associated with HOT regions show their stable expression pattern across all 35 analysed cell lines (Figure 1D), gene expression levels for those genes are generally above the median level of expression for all genes in the respective cell types. Gene Ontology (GO) [14] analysis revealed a variety of biological processes highly represented in HOT region-associated genes such as RNA processing, ncRNA processing, ncRNA metabolic process and ribosome biogenesis (Figure 1E), which is in line with the findings reported by [3].

To summarize, consistent with the published features of HOT regions [3,4], our findings confirm that genes associated with HOT regions are mostly housekeeping genes - they are required for the maintenance of basic cellular functions and are constitutively expressed.

### HOT regions have specific k-mer content compared to control regions

We analysed sequence characteristics of HOT regions in human, mouse, *C.elegans* and *D. melanogaster*, since sequence characteristics may be shared across species and that could explain the existence of HOT regions on multiple species. For this purpose, we built machine-learning models that can discriminate HOT regions from non-HOT or so called “COLD” regions using sequence features. The machine-learning model is primarily used for identifying features that are predictive of HOT regions. For this purpose, we used low-level sequence features such as k-mers. We used 2bp, 3bp and 4bp long k-mer frequencies. In addition, we used GC content and CpG observed/expected ratio (O/E ratio). CpG islands are a frequent feature of HOT regions in human and mouse. Although, *C. elegans* and *D. melanogaster* do not have CpG islands, CpG enrichment could be important at least for *C.elegans*, for which HOT regions are enriched for CpG dinucleotides [10]. We built a predictive model of “hotness” of genomic regions using a penalized multivariate regression method [15]. We built four species-specific models using normalized feature matrix as input. We had high accuracy for all models: cross-validation AUC between 0.82 and 0.94 for all the models. The top 10 feature importance averaged across species shows that CpG and GC rich k-mers along with CpG O/E ratio are the most important predictors for all the models (See Figure 2A for feature importance across all models and Figure S1A for individual models). The most predictive features averaged from all species are sufficient for discriminating HOT and COLD regions for all species. Although, localized CpG and GC spikes across genomes of *C. elegans* and *D. melanogaster* are not common, we can discriminate HOT and COLD regions across all four species using the same GC/CpG rich top features. We used principal component analysis to visualize the discrimination between HOT and COLD regions using the top features (Figure 2B). To determine whether there are higher order sequences which differentiate between HOT and non-HOT regions, we performed discriminative *de novo* motif analysis on HOT regions from all four species. The resulting motifs were short (5-6 bp with high information content), GC and CpG dominant (Figure S1B). The motifs partially matched binding sites of known transcription factors which bind GC rich sequences, such as SP1.

**Figure 2.**
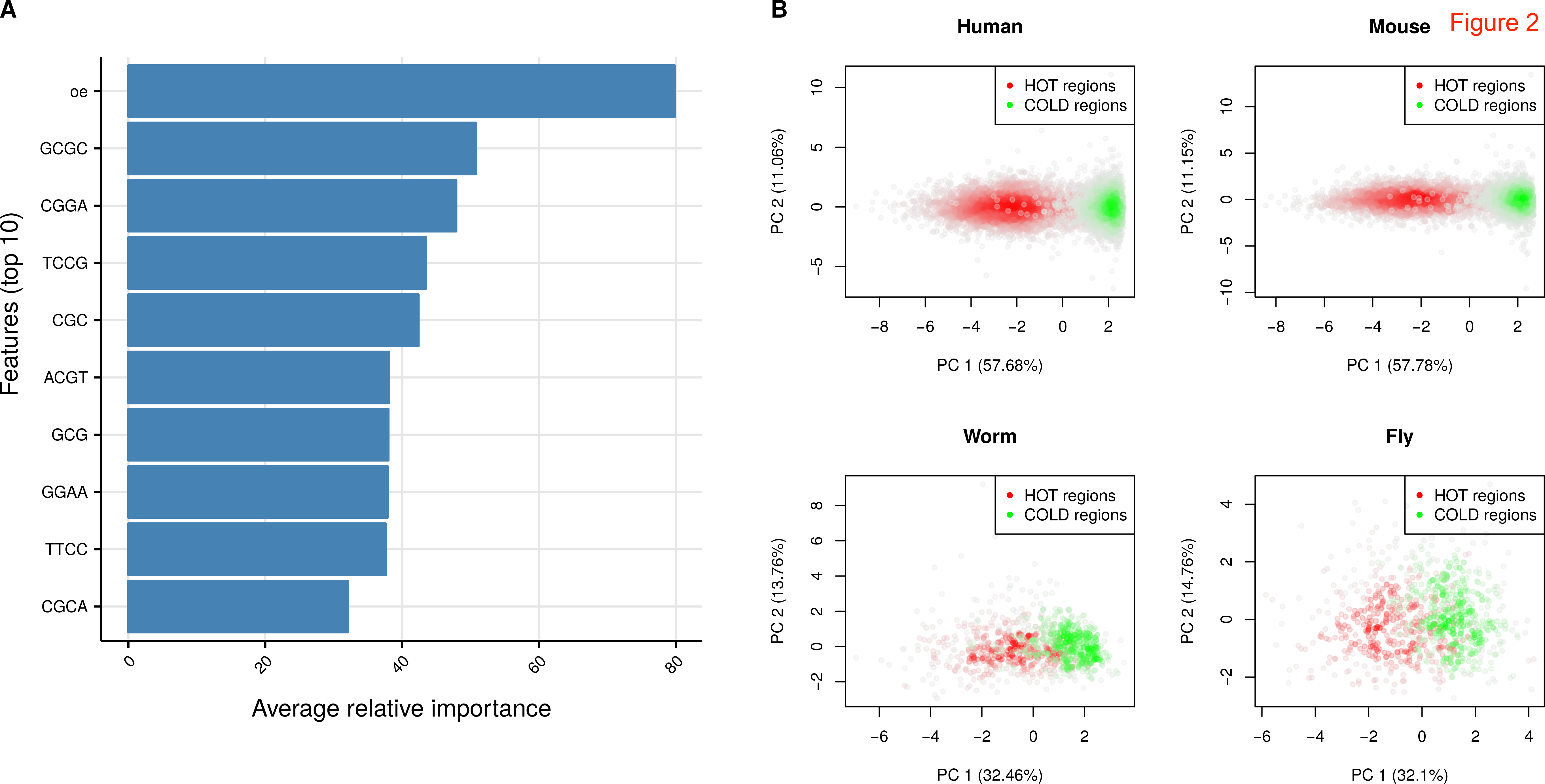
K-mer properties of HOT regions. A) Top 10 features ordered in relative importance averaged across species. Importance scores are scaled to 0-100 scale for each species then averaged. B) Principal component analysis using top 10 features shown in A. PCA is carried out for human, mouse, worm and fly separately. Scatter plots using first two principal components are shown, each dot represent HOT and COLD regions. The scatter plot is colored based on density of points, the more dense the points the darker the color.

### ChIP-seq for knock-out transcription factors have enrichment in HOT regions

Upon observing common low-level sequence features of HOT regions across species, we investigated whether potential technical biases of the ChIP-seq method could at least partially explain false positive signals on HOT regions. Previous studies suggest that even if the ChIP-ed protein does not exist in the analysed sample, highly expressed loci might give rise to false-positive peaks in yeast [7] and *D. melanogaster* [11]. In order to address this question with a more comprehensive collection of datasets, we downloaded all available experiments where the ChIP-ed transcription factor was not expected to be physically present in the cell as the gene encoding the transcription factor was “knocked-out”. This set consists of 43 ChIP-seq experiments for knock-out (KO) transcription factors (KO ChIP-seq), where only 24 experiments have a control experiment in the form of input DNA or mock-IP (See Supplementary Table 1 for accession numbers and details). These experiments are carried out by different labs, which reduces the lab-specific bias for KO generation and ChIP-seq experiments. More than half of the KO ChIP-seq experiments show a clear signal enrichment (measured as IP/control) over HOT regions. KO ChIP-seq experiments with strong enrichment on HOT regions are shown in Figure 3A and experiments without signal enrichment are shown in Figure S2A. The signal is absent from regions which do not have extreme enrichment of TF binding events. Pooling all the available signal enrichment for the KO ChIP-seq experiments with strong enrichment on HOT regions also shows the trend where signal enrichment on average is higher for HOT regions (Figure 3B shows signal enrichment of HOT regions and other control regions binned based on their TF occupancy percentiles).

**Figure 3.**
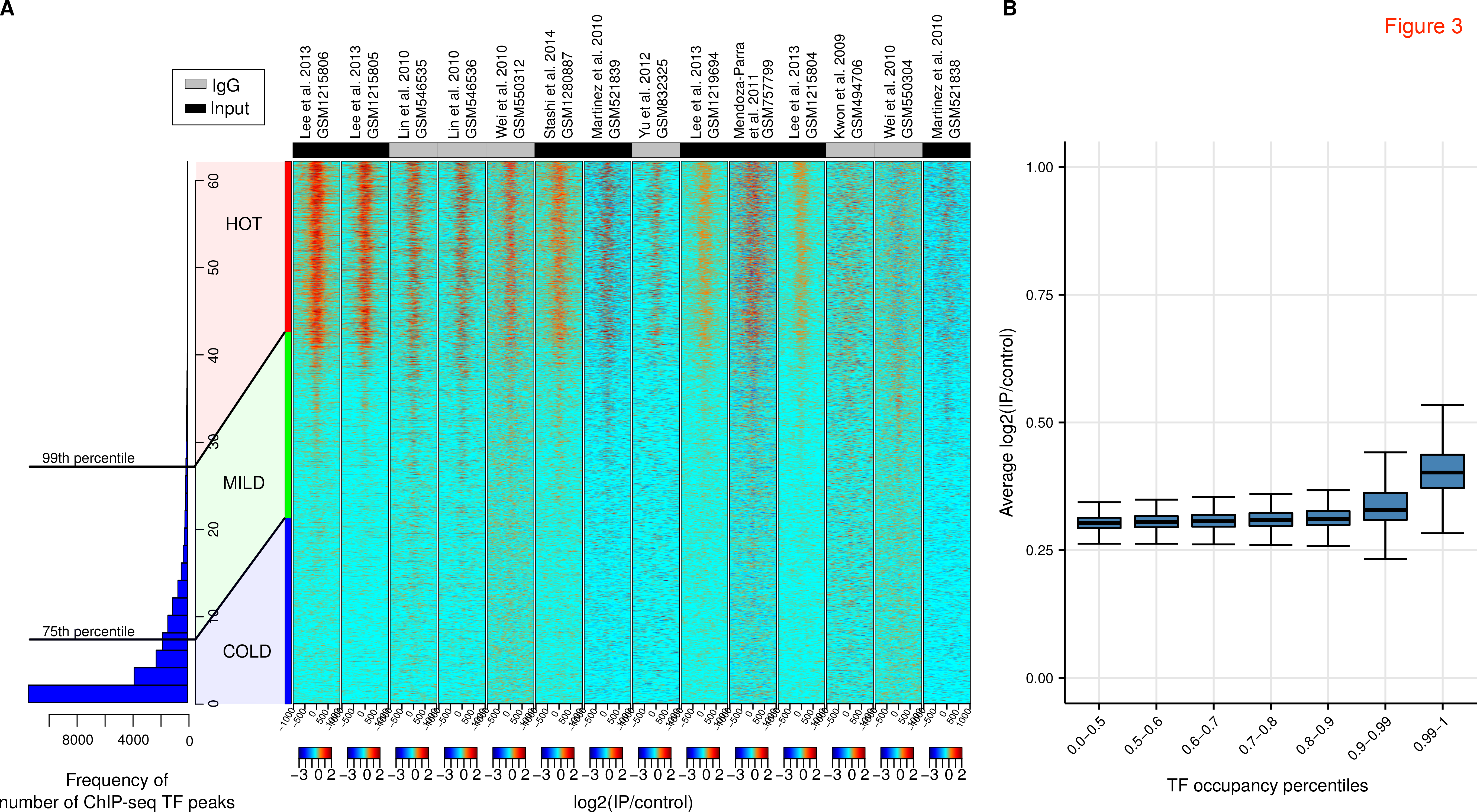
A) Heatmaps show KO ChIP-seq experiments with signal on HOT regions. The barplot indicates number of ChIP-seq peaks are in HOT (red), MILD (green) and COLD (blue) regions. The values in the heatmaps are log2(IP/control). The status of the control sample input DNA or IgG is color coded on the top of the heatmaps. B) Boxplots show distribution of average log2(IP/control) values for different sets of TF occupancy percentile bins. The log2(IP/control) values for each region in A is averaged across KO ChIP-seq experiments. The rightmost boxplot represents HOT regions, TF occupancy > 99th percentile, and the rest represent control region sets with different TF occupancy percentiles.

In addition, we noticed that some KO ChIP-seq experiments used IgG “mock” ChIP-seq as control. The IgG ChIP-seq experiments should ideally control for unspecific binding that could potentially cause a false positive signal, and yet more than half of KO ChIP-seq experiments that have IgG ChIP-seq as control show signal enrichment on HOT regions (see Figure 3A). Following up on this, we wanted to see whether the HOT regions show an enrichment of signal in IgG control experiments. We downloaded available IgG control experiments from ENCODE, where antibodies from the same vendor was used in multiple cell types (results shown in Figure S2B). HOT regions showed a consistent enrichment in multiple IgG experiments, however, the enrichment was weak and showed variability, which was dependent on the cell type (Figure S2C).

### HOT regions are associated with R-loops and G-quadruplex DNA

We next investigated the association of HOT regions with other GC rich features of the genome. One such feature that shares the same type of annotation with HOT regions such as CpG islands is R-loops. An R-loop is a nucleic acid structure that is composed of an RNA-DNA hybrid and consequently displaced single-stranded DNA [16]. Their formation and stabilisation are associated with GC content and CpG islands [17] and G-quadruplexes [18]. R-loops exist across a broad spectrum of species from bacteria to high eukaryotes [19] and are shared across mammals [19,20]. R-loop accumulation is a source of replication stress, genome instability, chromatin alterations, or gene silencing. They are associated with cancer and a number of genetic diseases [16].

R-loops can be detected genome-wide using a method called RNA-DNA immunoprecipitation followed by sequencing (DRIP-seq). It involves immunoprecipitation and sequencing of DNA fragments using the RNA-DNA hybrid specific S9.6 antibody [21], which was developed by extensively testing for specificity to RNA-DNA hybrids [22]. We analysed publicly available DRIP-seq datasets to investigate R-loop enrichment on HOT regions [20,23,24] (See Supplementary Table 1 for accession numbers). We observed R-loop enrichment on HOT regions on every cell line analyzed compared to other region sets binned based on their TF occupancy percentiles (Figure 4A). We observed this enrichment even when the DRIP-seq experiments with RNAseH treatment were used as controls. The RNAseH treatment removes R-loops and subsequent DRIP-seq experiment results in depleted signal for R-loops. This shows that the S9.6 antibody binds specifically to R-loops and does not show additional interactions with other forms of DNA and DNA-binding proteins. In addition, we also observed DRIP-seq enrichment on HOT regions of *C. elegans* (Figure 4B). These results suggest that R-loops across different species overlap with HOT regions.

**Figure 4.**
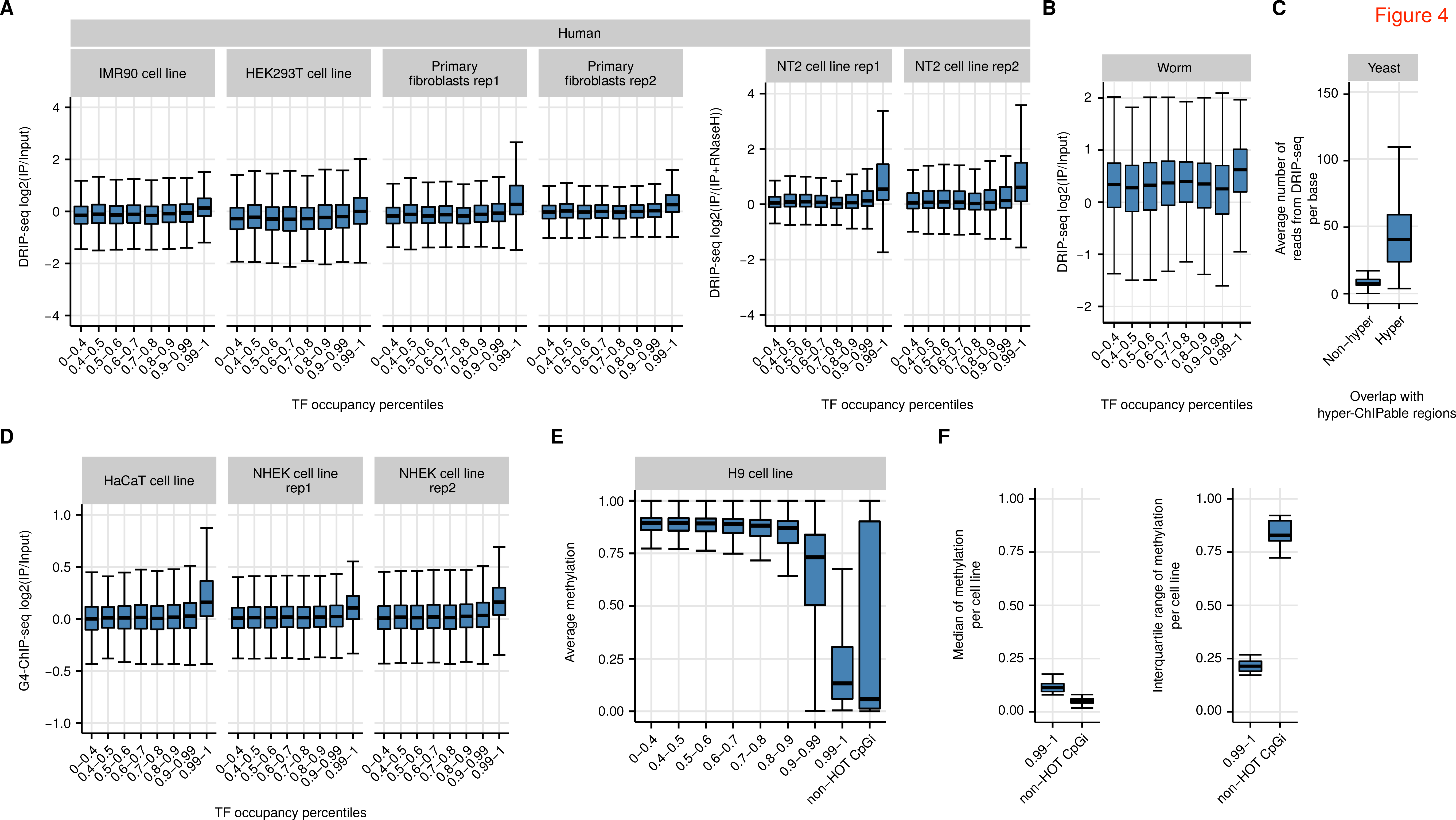
R-loops are associated with HOT regions. The boxplots show log2(IP/control) for HOT regions and control regions binned based on their TF occupancy percentile A) Various human cell lines and B) in worm. Boxplots show DRIP-seq read count per base-pair for hyper-ChIPable regions and all other genes as controls. D) HOT regions are enriched with G-quadruplex DNA (G4-ChIP-seq). Boxplots show log2(IP/control) for HOT regions and control regions binned based on their TF occupancy percentile. E) HOT regions are hypo-methylated in comparison to controls in H9 cell line. Boxplots show distributions of methylation for HOT regions (rightmost boxplot) and control regions binned based on their TF occupancy percentile F) Left Boxplot showing distributions of methylation medians across cell types for HOT regions and CpG islands that are not associated with HOT regions (non-HOT CpGi). Right boxplot showing distributions of methylation IQRs (interquartile ranges) across cell types for HOT regions and non-HOT CpGi.

R-loops usually colocalize with G-quadruplex DNA (G4-DNA) which is a tertiary structure of single-stranded DNA [16]. These structures can form on the displaced single-stranded G-rich DNA on the opposite side of the R-loop. We calculated the enrichment of G4-DNA on HOT regions using G4-ChIP-seq experiments [25], which are shown to enrich for G4-DNA specifically. We observed enrichment of G4-DNA signals on HOT regions, which is consistent with R loop localization on HOT regions (Figure 4D).

In addition, if R-loops are associated with hyper-ChIPability that gave rise to the definition of HOT regions, we would also expect to see R-loops in hyper-ChIPable regions originally defined in yeast by Teytelman et al. [7]. Indeed, we see enrichment of DRIP-seq signal [26] on published hyper-ChIPable regions of yeast (Figure 4C).

Since R-loops are associated with HOT regions, occupying TFs must be able to bind RNA-DNA hybrids or single-stranded DNA (ssDNA). Therefore, we checked if the TFs assayed by ENCODE have ssDNA binding or RNA-DNA hybrid binding domains, or such GO term annotations. Out of 165 studied TFs in human, only 2 of them contain at least one of the ssDNA binding domains: BACH1 contains a ‘DUF1866’ domain (from the RRM clan) and E2F6 contains a BRCA-2_OB3 domain (from the OB clan). Furthermore, none of the 165 TFs have an annotation of the GO term ‘single-stranded DNA binding’ (GO:0003697). When considering the direct interaction partners of these TFs, 31 out of 165 TFs (18.8%) have at least one direct interaction partner with an ssDNA-binding domain and 11 out of 165 TFs (6.7%) have at least one direct interaction partner with the GO term annotation for ssDNA binding. On the other hand, Cauli_VI domain that mediates the binding of RNASEH1 to RNA/DNA hybrids, is annotated only for two proteins in the whole proteome (RNASEH1 and Ankyrin repeat and LEM domain-containing protein 2 (ANKLE2)) and none of the human proteins have the associated GO term ‘DNA/RNA hybrid binding (GO:0071667)’ (according to the reviewed UniProt sequence annotations). Therefore, we could not detect any association of TFs or TFs’ interaction partners with RNA/DNA hybrid binding function.

### HOT regions have stable hypo-methylation across cell types

We also investigated the CpG methylation dynamics over HOT regions using base-pair resolution methylation data across multiple human cell types. Since most HOT regions are associated with CpG islands and above average expressed genes, we would expect low methylation over HOT regions [27]. In addition, hypo-methylated CpGs are prevalent in R-loops, and the formation of R-loops are proposed to be protecting the R-loop associated loci from de novo DNA methylation [16]. Consistent with these information, we observed hypo-methylation for HOT regions compared to controls. The median methylation levels for HOT regions was similar to the median methylation levels of CpG islands not associated with HOT regions (non-HOT CGI) (See Figure 4E for an example cell line, see Figure S3 for all analyzed cell lines). Interestingly, non-HOT CGI had higher variation of methylation than the HOT regions despite the median methylation for both sets being low. This was a trend evident in all the cell types examined. Across the cell types, non-HOT CGI had 3-4 times higher methylation variation than HOT regions. This indicates that although HOT regions are associated with CpG islands, they are different from non-HOT CpG islands in their methylation dynamics and they maintain low levels of methylation across different cell types (See Figure 4F).

## Discussion

HOT regions are locations in the genome with remarkably high occupancy of transcription factors. These regions are mostly associated with promoters of stably expressed genes. We showed that the low-level sequence features, such as GC rich and CpG containing k-mers, are shared across HOT regions of different species. Most interestingly, we demonstrated that HOT regions are specifically enriched with false positive signals, using KO transcription factor ChIP-seq. These false positive signals are antibody dependent since KO ChIP-seq experiments show variable intensity of signals on HOT regions. The traditionally suggested controls, such as IgG ChIP-seq, can not reliably control for these artifacts. We showed that HOT regions associate with R-loops, in multiple organisms, as well as G-quadruplex DNA structures. Our results support the view that the peaks observed on HOT regions are produced by the unspecific enrichment in multiple ChIP-seq experiments, rather than by the pull-down of specific transcription factors.

There might be many causes for the persistent false positive signal on HOT regions. The ChIP-seq signal consists of the signal from actual binding events and the noise. The noise is usually attributed to sequencing depth, library preparation, but most importantly to antibody specificity [28]. The observed false positive signal could be obtained through pull-down of non-target proteins; this would however require that all experimentally used antibodies cross-react with a small set of proteins which constitutively bind GC rich promoters in multiple cell lines - a scenario which is highly improbable. The degree of overlap of HOT regions with R-loops suggests another hypothesis - that the antibodies cross-react directly with polynucleotide epitopes present in the HOT regions [29]. R-loops are formed during transcription of GC rich, hypomethylated regions, where the nascent RNA strand displaces one of the DNA strands, forming an RNA:DNA Watson-Crick base pairing with the complementary strand. Such displacement causes R-loop prone regions to contain multiple polynucleotide structures: double stranded DNA, single stranded DNA, RNA:DNA hybrids, single stranded RNA (reviewed in [16]), G quadruplex complexes [30,31], etc., all of which can be bound by antibodies with a range of affinities [30,32–35]. Anti-DNA antibodies are abundant in the serum of normal animals immunized with protein fragments [36–45], and are frequently polyspecific [40,46–54] - they can bind both polynucleotide and non-polynucleotide (e.g. peptide, phospholipid) epitopes [35]. Abundance of epitopes in constrained genomic regions, along with the fact that the HOT regions are associated with CpG islands of housekeeping genes (which are expressed and form R-loops in many cellular systems), and the promiscuity of antibodies, provide a simple explanation for the ubiquity of enrichments observed on HOT regions in various ChIP-seq experiments. Serum of non-immunized, healthy animals usually contains a low percentage of anti-DNA binding antibodies. This explains why the IgG samples, when used as controls, show a signal on HOT regions, but the intensity of the signal is much lower than from antibodies produced by deliberate immunization. The recommended experimental methods for ascertaining antibody specificity [55] control almost exclusively for binding of antibodies to non-target proteins, so the direct interaction of antibodies with polynucleotide epitopes might be an underappreciated source of false positives in ChIP-seq experiments (Our model summarized in Figure 5). The signal on HOT regions could additionally arise by direct binding of TFs to single-stranded DNA (ssDNA) or RNA-DNA hybrids. Based on the current protein domain annotations, few to none of the TFs have such capabilities.

**Figure 5.**
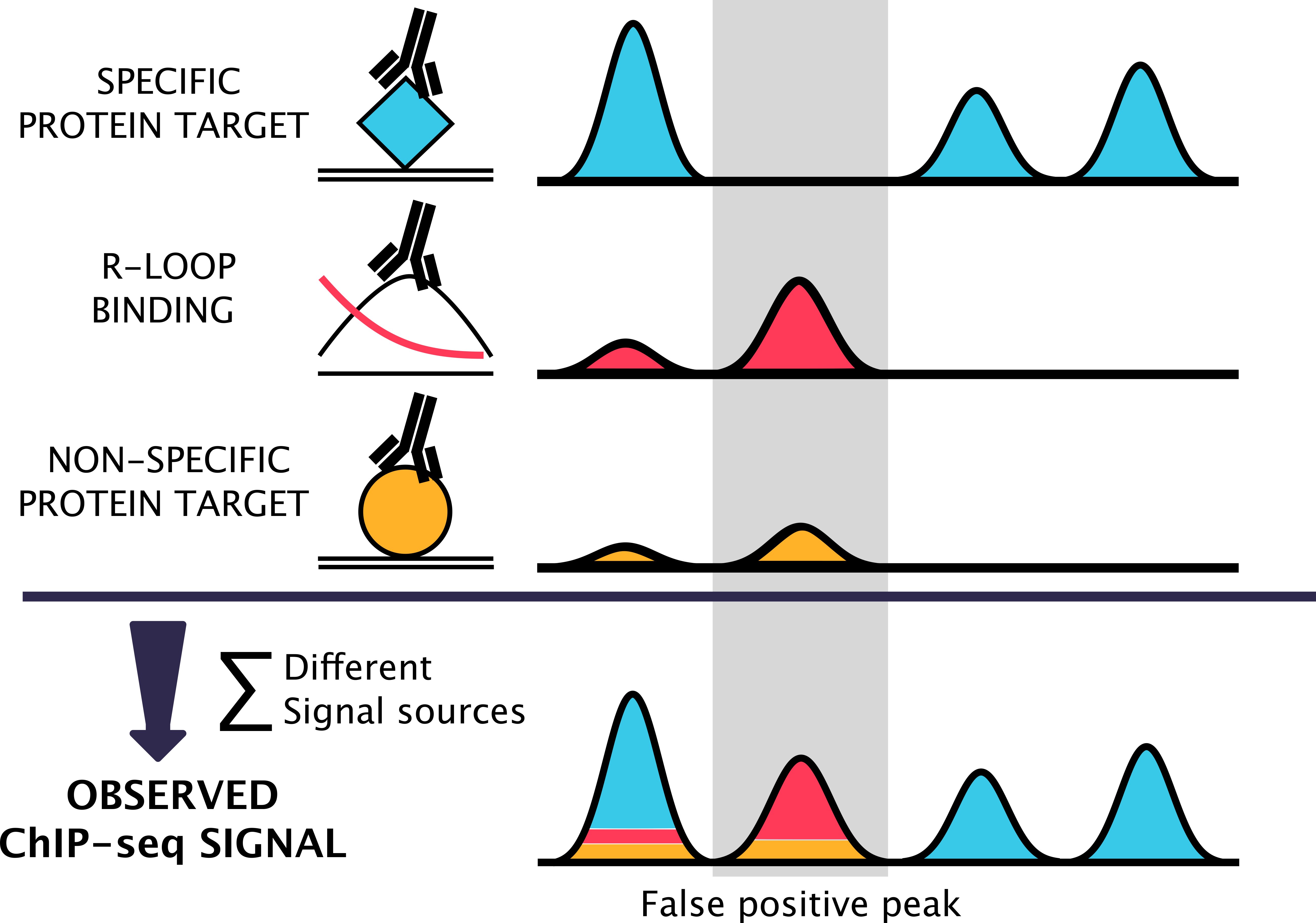
The observed ChIP signal arises from a combination of different signal sources. The signal in a ChIP experiment originates from an antibody binding to the intended target protein (blue), and nonspecific antibody binding - either to the non target proteins (orange) or directly to polynucleotide structures, such as R-loops (red). The error (orange + red) is not proportional to the signal from the targeted protein, rather, it depends on sequence properties and expression characteristics of individual genomic regions. This results with a subset of enriched regions in the ChIP experiments being false positives.

In this work, we have focused on regions that show high enrichment in multiple ChIP-seq experiments. Although we provide evidence that HOT regions do not contain several dozens of bound transcription factors, the real extent of detected false positive interactions is probably not limited to HOT regions. With the currently available data, it is not possible to estimate the proportion of an antibody specific error resulting from the enrichment due to the pull-down of non-target proteins vs. the direct binding to polynucleotide epitopes. Examination of the DNA binding properties of monoclonal antibodies, for example with protein binding arrays [56,57], might provide the required data for constructing more precise error models.

Lack of a strong signal over HOT regions, in a subset of KO ChIP-seq samples shows that by using stringent antibody validation methods, it is possible to perform highly specific ChIP experiments. A level of prudence is needed though - a lack of signal in a KO ChIP-seq experiment might also be caused by technical conditions such as low number of reads, low library complexity or unsuccessful IP.

Our results, yet again [29,55,58,59], emphasise the need for critical examination and extensive testing of antibodies prior to their experimental usage. Whenever possible, controls in ChIP-seq experiments should be performed by ChIP-ing of protein in a system where the protein is not physically present, as implemented in Knockout Implemented Normalization method (KOIN) [60]. If such controls are unfeasible, we provide lists of HOT regions and ChIP-seq peaks overlapping with those regions should be carefully examined. Either those peaks should be removed from the set or more stringent quality controls, such as requirement of a canonical motif, should be exercised before making correlative or functional conclusions about the protein of interest.

## Methods

### ChIP-seq data for definition of HOT regions

Analyses of human TF binding sites were performed using the UCSC human hg19 reference genome, mouse mm9, *D. melanogaster* dm3 and *C. elegans* ce10. ChIP-seq files in narrowPeak format were downloaded from the ENCODE (www.encodeproject.org) and modENCODE (data.modencode.org) portals. Human TF binding sites in narrowPeak format were downloaded from the UCSC Uniform track http://hgdownload.cse.ucsc.edu/goldenPath/hg19/encodeDCC/wgEncodeAwgTfbsUniform/. 166 human TFs, 42 murine, 42 *D. melanogaster*, and 83 *C. elegans* were obtained.

### Defining HOT regions

For a given set of ChIP-seq peaks per species, we determined the summits of the peaks. Following that, we calculated the density of the summits over the genome using 500 bp sliding windows. We calculated the local maxima of the density vector for each chromosome. We made sure local maxima of the density vector are the only maxima in 2000 bp surrounding of the maxima for human and 1000 bp for other species. This is necessary to remove sub-optimal maxima around the real maxima. 2000 bp threshold was specifically applied for human datasets due to high number of experiments creating multiple local maxima around the real maxima. We then ranked these maxima based on the density scores, which is effectively the number of overlapping ChIP-seq peaks and represents the TF occupancy. These density scores are referred to as TF occupancy throughout the text. We used 99th percentile threshold to define the HOT regions. This is in line with previous methods [61]. HOT regions were called using only the regulatory peak sets (no RNA polymerase datasets were included). The regions that are not selected as HOT regions are binned according to their TF occupancy percentiles (number of ChIP-seq peak counts) and used as controls in follow-up analyses.

### Assigning HOT regions to genes and expression analysis

Distance from HOT regions to the nearest transcription start sites was analysed using GREAT [14]. Expression values of genes associated with HOT regions across tissues were obtained from the Expression Atlas EBI database (www.ebi.ac.uk/gxa) and fantom5 CAGE expression [62].

### Sequence analyses of HOT regions

Extraction of 2-,3 and 4-mers, CpG frequencies (sum of observed G and C divided by length of genomic region equal to 2000), GC skew, the observed/expected ratios for CpG on HOT regions was computed using scripts written in R version 3.3.1 and BSgenome package. We used the following genome assemblies: mm9, hg19, ce10, dm3. The observed/expected ratios for CpG were calculated according to the formula (O/E)_CpG_ = [f(CG)/f(C)f(G)] × width(genomic_region), where f denotes the observed frequency of the given mono- or di-nucleotides. De novo motifs were found using R package motifRG [63].

MotifRG is a discriminative motif analysis tool which searches for overrepresented motifs in a positive set when compared to a negative set. Sequences from HOT regions were used as the positive set, while the negative set was constructed by sampling equal number of sequences as there are HOT regions from non-HOT regions. Motif analysis was performed on separately on HOT regions from *H.sapiens*, *M.musculus*, *D.melanogaster* and *C.elegans*. MotifRG was run with the following parameters: start.width=6, both.strand=TRUE, mask=TRUE, enriched.only=TRUE.

### Elastic net construction and PCA for discrimination of HOT regions

For training, we used HOT regions defined as regions with TF occupancy percentiles higher than 0.995 for hg19 and 0.99 for other organisms. In order to use as a control, regions with TF occupancy lower than 85th percentile, were sampled matching the number of selected HOT regions. HOT and control regions for mm9 and hg19 were CpG sampled, in order to sure that the ratio of HOT and control regions that overlap CpG islands were the same. The genomic coordinates of the CpG islands were downloaded from the UCSC Table browser (cpgIslandsExt table). All models had the following set of features: CpG frequencies, ratio of observed versus expected CpGs, GC skew, and 2 3-, 4-mers. Feature matrix was standardized prior to training.

Models trained and tested for the same species were trained using 10-fold cross-validation. Variable importance scores were calculated for each species-specific model as an absolute value of the model coefficients, which were then normalized to a scale from 0 to 100. Average relative importance was calculated as an average of variable importance scores of all models. Area under the ROC curve (AUC) was computed to measure the accuracy of the models. We used elastic net function from the glmnet R package [64]. For the PCA, we used top 10 features ranked by the average relative importance. Using these same features for all species, we calculated PCA and plotted the the color coded scatter plot on principal components for each species. For illustration purposes, we sampled the same number of “COLD” regions as the number “HOT” regions.

### Data processing and visualization of KO ChIP-seq, DRIP/RDIP-seq and G4 ChIP–seq samples

Fastq files of KO ChIP-seq experiments (See Supplementary Table 1) were downloaded from the European Nucleotide Archive database (ENA). All fastq files have single-end reads and were uniquely mapped into murine genome version mm9 using Bowtie 1.1.12 with parameters: -p 3 -S -k 1 -m 1 –tryhard -I 50 -X 650 –best –strata –chunkmbs 1000. The bbduk program from the BBMap software 35.14 was used for adapter, quality trimming and filtering with parameters: minlength=20 qtrim=r trimq=20 ktrim=r k=25 mink=11 ref=”bbmap/resources/truseq.fa.gz” hdist=1. Two of KO ChIP-seq samples NFAT1_P+I and NFAT1_None didn’t pass bbduk tests and were excluded from the further analysis. The FastQC 0.11.3 program was used for quality control. Conversion from SAM to BAM file format, sorting and indexing BAM files was done using samtools 0.1.19, conversion from BAM to BED file formats and then BED to BedGraph file formats using Bedtools-2.17.0, from BedGraph to BigWig file format using BedGraphToBigWig v4. The same pipeline was used for DRIP-seq and RDIP-seq samples, and G4 ChIP-seq (due to lack of detected adapters, bbduk argument ‘ref’ that indicates a path to adapters was omitted).

The R package genomation [65] was used for calculating fold enrichment analysis of KO ChIP-seq samples and plotting heatmaps. Fold enrichment of KO ChIP-seq samples was defined as log2 of IP signal divided by control per base pair. Out of total 25 KO ChIP-seq samples, 15 are positively and 9 are negatively associated with TF occupancy scores. Heatmaps were binned on x-axis into 50 bins, average for each bin was taken, and winsorized to limit extreme values below 0.5 and above .99 percentile. Some of KO ChIP-seq samples are conditional knockouts using Cre-lox recombination system (See Supplementary Table 1). Enrichment presented as boxplots of KO ChIP-seq, DRIP/RDIP-seq, G4-ChIP-seq samples (Figure 3B and 4A,B,D) was calculated as the logarithm base 2 of the number of reads from IP sample overlapping HOT regions (normalized for library size and multiplied by counts per million) divided by the number of the reads from the control sample overlapping HOT regions normalized in the same way. If control was not available then IP with RNaseH treatment was treated as control. For visualisation purposes, windows on heatmaps were sampled: 3000 windows from 0 to 75th TF occupancy percentile, 3000 from 75th to 99th, and 3000 from 99th to 100th percentile.

IgG samples corresponding to the following antibody ENCAB000AOJ were downloaded from ENCODE. Samples marked with “extremely low read depth were removed from the analysis”. Samples which belong to the same biosample term id were pooled together. Signal was visualized in a region of +/- 1kb around HOT (>99th percentile), MILD (between 99th and 75th percentile), and COLD regions (below 75th percentile). Prior to visualization, the reads were extended to 200bp in a stranded fashion, and the signal was normalized to per million reads.

### Methylation dynamics for HOT regions

Methylation over the regions of interest is extracted from the Roadmap Epigenomics Consortium Whole-genome Bisulfite sequencing data sets [66]. Our regions of interest consist of HOT regions, non-HOT regions (regions with lower TF occupancy), and CpG islands not associated with HOT regions (non-HOT CGI). For each region of interest, we extracted overlapping methylation value for each cell type and calculated the mean methylation value per region. We plotted the distribution of mean methylation values for each set of regions: HOT regions, non-HOT regions (binned into different TF occupancy levels), and non-HOT CGI. For each cell type, we calculated the interquartile range and median methylation values for HOT regions and non-HOT CGI. Next, we plotted the distributions of medians and interquartile ranges across cell types as boxplots to compare the methylation dynamics for HOT regions to non-HOT CGI.

### PFAM domains and Human Protein-Protein Interactions

Reviewed UniProt [67] human protein sequences (as of 31.08.2015) were scanned for occurrence of PFAM HMM (both PFAM-A and PFAM-B) models using HMMER3 [68]. The HMM scanning detected 9511 types of PFAM domains in 19275 proteins.

PFAM [69]entries for single-stranded DNA-binding domains were collected from the PFAM database by combining all members of the following PFAM clans: OB (for OB-fold domains), KH (for K-homology domains), RRM (for RRM-like domains), and sPC4-like (for the Whirly domain). The collection of these four clans contains 90 different types of PFAM domains.

Human protein-protein interaction data was downloaded from the iRefWeb database [70]. In order to dissect which protein-protein interactions of TFs are direct (or relatively direct physical interactions of proteins), the interactions were filtered for the following criteria: 1)Interactor A (uidA) is from taxa:9606 and interactor B (uidB) is from taxa:9606; 2)Interaction type between uidA and uidB is one of ‘MI:0915(physical association)’, ‘MI:0407(direct interaction)’, ‘MI:0403(colocalization)’, ‘MI:0914(association)’, or ‘MI:0191(aggregation)’; 3)Both uidA and uidB have an ID mapped to UniProt accessions.

## Acknowledgements

We are grateful for the core funding from Helmholtz Association. In addition, BU was supported by the RNA Bioinformatics Center of the German Network for Bioinformatics Infrastructure (de.NBI) [031 A538C RBC (de.NBI)]. KW is supported by the Berlin Institute of Health.

## Contributions

AA and VF conceived the idea during discussions on ChIP-seq noise on ENCODE datasets. AA designed the study with input from VF and KW. KW downloaded and processed all the ChIP-seq, DRIP-seq, G4-ChIP-seq data from human, mouse, worm and fly. Yeast DRIP-seq data is processed by VF. HOT region algorithm is designed and implemented by AA with contributions from KW. HOT region predictions for each species is done by KW. Machine-learning approach is implemented by AA, KW and VF. BU and RW provided support with data analysis, processing for peak calling and examining HOT region sequence characteristics. KW, VF, AA and BU wrote the manuscript. AA supervised the project and ensured its progress.

